# Heterogenous susceptibility to R-pyocins in populations of *Pseudomonas aeruginosa* sourced from cystic fibrosis lungs

**DOI:** 10.1101/2020.08.05.238956

**Authors:** Madeline Mei, Jacob Thomas, Stephen P. Diggle

**Author notes:** Address correspondence to Stephen P. Diggle. **Author contributions**: M.M., J.T. and S.P.D. conceived the study and wrote the paper; M.M. performed the experimental work and analyzed the data.

## Abstract

Bacteriocins are proteinaceous antimicrobials produced by bacteria which are active against other strains of the same species. R-type pyocins are phage tail-like bacteriocins produced by *Pseudomonas aeruginosa*. Due to their anti-pseudomonal activity, R-pyocins have potential as therapeutics in infection. *P. aeruginosa* is a Gram-negative opportunistic pathogen and is particularly problematic for individuals with cystic fibrosis (CF). *P. aeruginosa* from CF lung infections develop increasing resistance to antibiotics, making new treatment approaches essential. *P. aeruginosa* populations become phenotypically and genotypically diverse during infection, however, little is known of the efficacy of R-pyocins against heterogeneous populations. R-pyocins vary by subtype (R1-R5), distinguished by binding to different residues on the lipopolysaccharide (LPS). Each type varies in killing spectrum, and each strain produces only one R-type. To evaluate the prevalence of different R-types, we screened *P. aeruginosa* strains from the International Pseudomonas Consortium Database (IPCD) and from our biobank of CF strains. We found that (i) R1-types were the most prevalent R-type among strains from respiratory sources; (ii) there is a large number of strains lacking R-pyocin genes, and (iii) isolates collected from the same patient have the same R-type. We then assessed the impact of intra-strain diversity on R-pyocin susceptibility and found a heterogenous response to R-pyocins within populations, likely due to differences in the LPS core. Our work reveals that heterogeneous populations of microbes exhibit variable susceptibility to R-pyocins and highlights that there is likely heterogeneity in response to other types of LPS-binding antimicrobials, including phage.

**Importance:** R-pyocins have potential as alternative therapeutics against *Pseudomonas aeruginosa* in chronic infection, however little is known about the efficacy of R-pyocins in heterogeneous bacterial populations. *P. aeruginosa* is known to become resistant to multiple antibiotics, as well as evolve phenotypic and genotypic diversity over time; thus it is particularly difficult to eradicate in chronic cystic fibrosis (CF) lung infections. In this study, we found that *P. aeruginosa* populations from CF lungs maintain the same R-pyocin genotype but exhibit heterogeneity in susceptibility to R-pyocins from other strains. Our findings suggest there is likely heterogeneity in response to other types of LPS-binding antimicrobials, such as phage, highlighting the necessity of further studying the potential of LPS-binding antimicrobial particles as alternative therapies in chronic infections.

## Introduction

Pyocins are narrow-spectrum antimicrobials, specifically produced by *Pseudomonas aeruginosa*, that have antimicrobial activity against members of the same species (1–4). *P. aeruginosa* produces three types of pyocin, referred to as S-pyocins, F-pyocins, and R-pyocins (1–3). The focus of this work is R-pyocins, which are narrow-spectrum, phage-tail-like bacteriocins that vary by subtype (types R1-R5) (2, 3, 5–11). Each *P. aeruginosa* strain uniquely produces only one of these R-pyocin types (2, 3). As the variable C-terminus region of the tail fibers of sub-types R2-R4 are approximately 98% similar in amino acid sequence, they are often grouped together under the R2 subtype (12–14). R-pyocin subtype and specificity are conferred by differences in the “foot” of the tail fiber (Fig. 1), which is believed to bind to specific residues on the lipopolysaccharide (LPS) decorating the outer membrane of Gram-negative bacteria (12–26). A strain may be susceptible or resistant to any variation of the different R-types (2, 3, 14, 25, 27–29).

**Figure 1.**
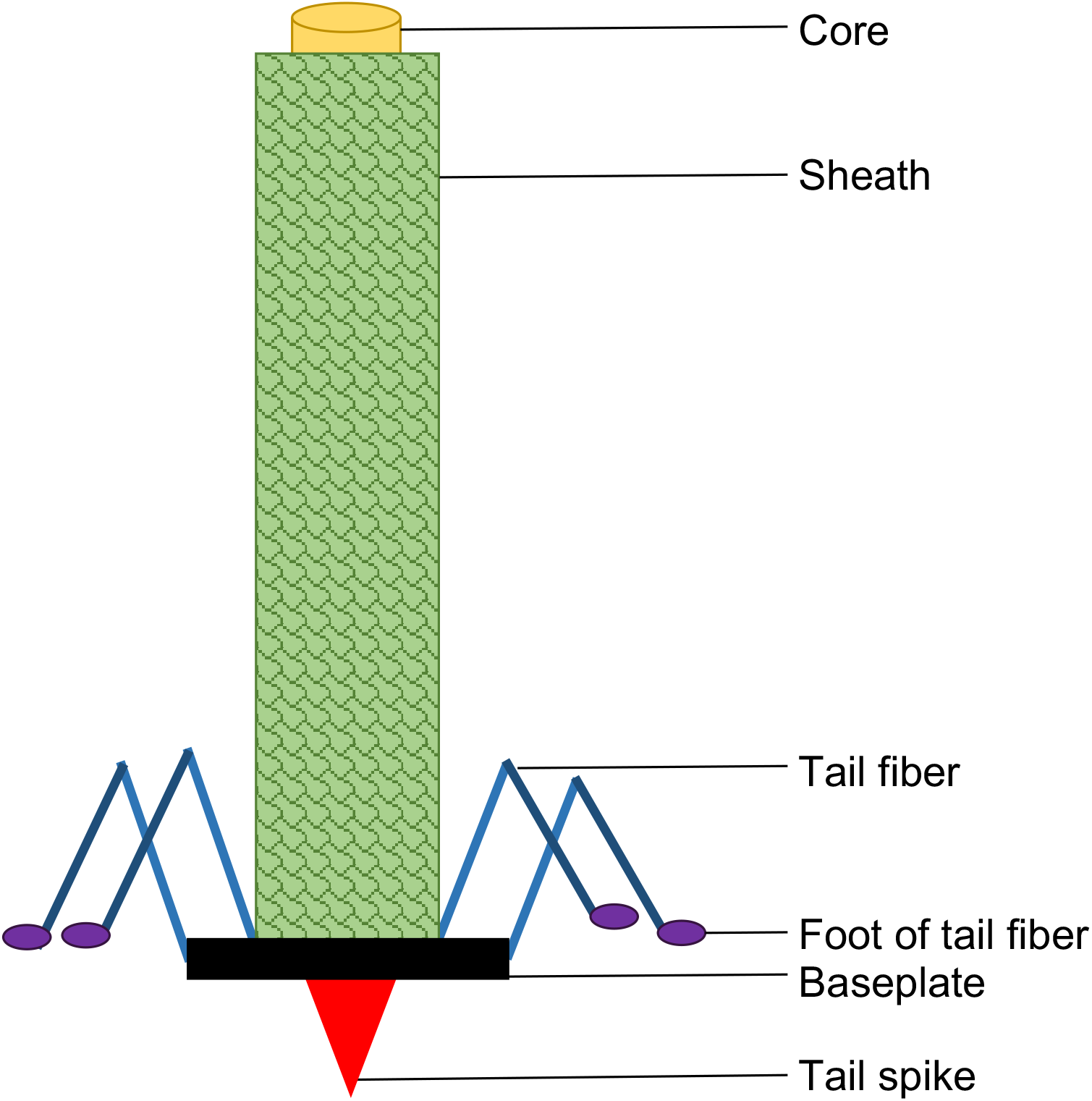
R-pyocin structure. R-type specificity is determined by the sequence on the foot of the tail fiber, which is believed to recognize glycosylated structures on the outer membrane (such as the LPS). Upon binding, the sheath contracts to push the core into the cell membrane. Puncture by the tail spike depolarizes the membrane potential, leading to cell death (1–3, 12, 14, 15, 22–27, 29–36). R-pyocin structure is depicted as described in previous studies (2, 3, 11, 14–21, 27, 29–31, 35, 36).

R-pyocins are similar to phage tails of the *Myoviridae* phage family, both in structure and in killing mechanism (2, 3, 14–17, 24, 27). The proposed mode of action is through the foot of the R-pyocin tail fiber recognizing a receptor on the LPS; binding to the target cell triggers the sheath to contract, pushing the tail spike and core through the cell envelope (1–3, 12, 14, 15, 22–27, 29–33). The puncture causes cytoplasmic membrane depolarization, inhibiting active transport and eventually leading to cell death (32–36). Several monosaccharide residues on the outer core of the LPS have been proposed to confer susceptibility and specificity to some of the R-pyocin types (24, 25), however, only the receptor of the R3-type pyocin has been clearly defined to be the Glc^IV^ terminal glucose on the outer core of the LPS (26). R-pyocins have previously been explored as potential therapeutics (12, 13, 37, 38), but little is known about their binding to target cells, or their killing efficacy in heterogeneous *P. aeruginosa* populations.

*P. aeruginosa* is found in a number of human infections and is the leading cause of morbidity and mortality in individuals with cystic fibrosis (CF). Once established in the CF lung, single-strain populations become heterogeneous - evolving genotypic and phenotypic diversity during the course of infection (39–49). *P. aeruginosa* possesses many intrinsic antibiotic resistance mechanisms, and isolates from CF lung infections are known to become increasingly resistant to multiple antibiotics over time (4, 42, 50–56). It is also known that isolates sourced from clinical populations of *P. aeruginosa* can exhibit considerable variation in susceptibility to antibiotics (44, 49, 54, 57–59), however heterogeneity in R-pyocin susceptibility, or heterogeneity in susceptibility to other types of bacteriocins have not been explored at the population level. Here we determined that R1-type pyocins were the most prevalent genotype among clinical respiratory strains (including CF), while a large proportion of strains lack R-pyocin genes entirely. We found that multiple isolates from diverse, clonal populations of *P. aeruginosa* have the same R-pyocin genotype and that R-pyocin type does not change across populations collected over a 2-year period within each patient. Moreover, we observed heterogeneity in R-pyocin susceptibility within a population of *P. aeruginosa in vitro.* This heterogenous response is likely due to differences in the LPS core composition among isolates, supporting evidence from other research groups that the LPS core contains the receptors for R-pyocins. Our findings highlight that diverse *P. aeruginosa* populations exhibit within-population heterogeneity in R-pyocin susceptibility, suggesting there is likely also heterogeneity in response to other types of LPS-binding alternative therapies, such as phage.

## Results

### R1 genotype is the most prevalent R-pyocin type in clinical *P. aeruginosa* strains

To determine the prevalence of different R-pyocin subtypes in *P. aeruginosa* strains sourced from CF infections, we conducted an *in silico* screen to determine R-pyocin genotypes of *P. aeruginosa* strains publicly available through the International *Pseudomonas* Consortium Database (IPCD) (60). The IPCD is a repository of *P. aeruginosa* strains (single isolates) collected from a variety of sources (with an emphasis on CF), created to facilitate metadata analyses specifically to improve prognostic approaches for CF treatment (60). By using the BLAST algorithm from NCBI, we screened 852 strains from the database for nucleotide sequence homology to R1-, R2-, or R5-pyocin tail fiber genes. As subtypes R2-R4 are highly similar, we considered these three subtypes as one larger R2-subtype for this analysis (12–15). Out of the 852 strains in the database, we could not conclusively type 303 strains. We then further screened the 303 untypeable strains for the presence or absence of genes flanking the R-pyocin gene cassette; these include trpE (PA0609), regulatory genes and holin (PA0610-PA0614), lytic genes (PA0629-PA0631), F-pyocin structural genes (PA0633-PA0648) and TrpG (PA0649). While all 303 strains contained trpE and trpG, regulatory genes (PA0610-PA0613), holin, and lytic genes, 297 strains were found to lack R-pyocin structural genes entirely. The six strains with intact and complete R-pyocin structural genes were confirmed to possess a full R-pyocin tail fiber gene, though it was not homologous to the known types (Supplementary Dataset 1). In addition, 300 of the untypeable strains still maintained some degree of F-pyocin structural genes. It is important to note that typing alone does not have implications on whether a strain produces functional R-pyocins or not.

Of the 852 strains in the database, we categorized 448 strains as isolated from human “respiratory” sources; these include *P. aeruginosa* isolated from throat, CF, bronchiectasis, sputum, sinus, and nasopharynx. Among the 448 respiratory strains, we typed 144 (32.14%) strains as R1-type, 76 (16.96%) as R2-type, and 43 (9.6%) as R5-type pyocin producers. The remaining 185 (41.29%) of strains were untypeable, with only two of these strains possessing R-pyocin structural genes. Fig. S1 depicts the distribution of R-types for each source; more information regarding the categorization and typing of strains from the IPCD can be found in Supplementary Dataset 1. Due to the curation and labeling of the strain information in the database, we were not able to precisely distinguish all CF strains from strains isolated from other respiratory infections, however is it likely that many of the samples labeled by anatomical respiratory sources are from CF patients, given the research areas of the curating research groups. The distribution of R-pyocin types among the typeable respiratory strains shows that (i) R1-pyocin producers are likely more prevalent in CF than the other R-pyocin subtypes (Fig. 2A) and (ii) a large proportion of strains lack R-pyocin structural genes entirely, though they maintain the flanking regulatory elements.

**Figure 2.**
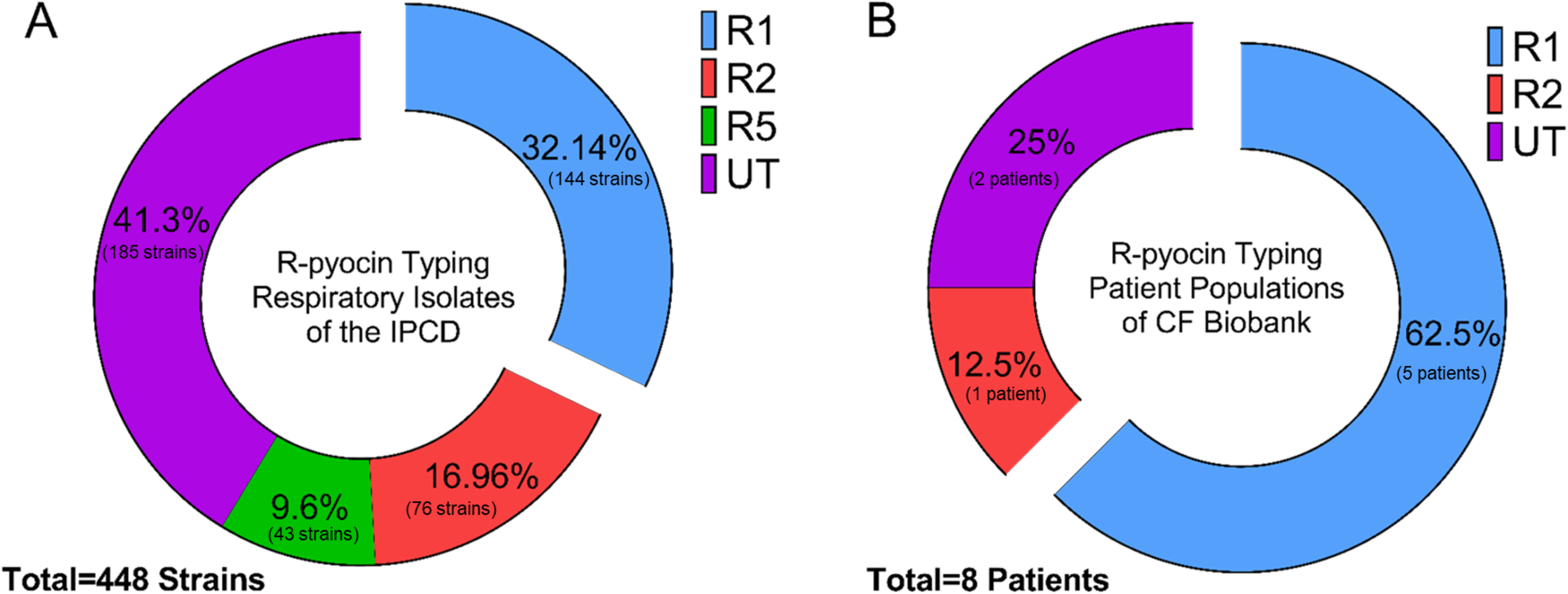
R-pyocin typing of clinical *P. aeruginosa* strains and whole populations isolated from CF patients. (A) Using BLAST homology-based alignment, we R-pyocin-typed publicly available *P. aeruginosa* sequences from respiratory sources of the IPCD to determine the prevalence of each pyocin type. Out of the 448 respiratory isolates in the entire database, 185 strains were not able to be typed. (B) Using multiplex PCR, we R-pyocin typed our own biobank of whole populations of *P. aeruginosa* isolated from expectorated sputum of CF patients to compare the prevalence of R-types found among CF patients in our cohort. We used 19 longitudinal samples collected from 8 patients and R-pyocin-typed 183 isolates total. We found that in our biobank, R1 was also the most prevalent R-pyocin type found among CF patients in our cohort (UT = Untyped R-pyocin genotype).

### R-pyocin type does not vary within clonal CF populations of *P. aeruginosa*

Through a collaboration with Children’s Healthcare of Atlanta and the CF Discovery Core at Emory University, we collected whole populations of *P. aeruginosa* from fresh CF sputum samples of patients with chronic lung infections. In this work, we use the term “population” to describe a collection of bacteria (*P. aeruginosa*) isolated from the expectorated sputum of a CF patient at a specific time point. Up to 10 isolates were chosen at random to R-pyocin type for each population, from each patient. Using multiplex PCR (Table S1), we R-pyocin typed isolates from 19 of these populations (183 isolates total) collected from 8 different patients over a 2 year period (Supplementary Dataset 2). In our cohort, the R1 genotype was the most prevalent R-pyocin type found among CF patients. We found that while R-pyocin type does not change within a population or over longitudinal collections from the same patient, different patients from the same clinic may be infected with *P. aeruginosa* containing different R-types. Out of the 8 patients we typed, 5 of these patients were chronically infected with an R1-pyocin producing strain, 2 patients were infected with untypeable populations, and 1 patient was infected with an R2-pyocin strain (Fig. 2B). It appeared that in the time frame sampled (over a 2-year period) there were no strain switching events within these patients. We did not find any populations containing the R5-pyocin type in our biobank.

### R-pyocin-mediated killing varies among heterogeneous *P. aeruginosa* populations

Previous work has shown that isolates from within the same population have diverse antibiotic resistance profiles (54), but diversity in R-pyocin susceptibility has yet to be evaluated within whole populations of *P. aeruginosa*. We chose 20 random isolates from three *P. aeruginosa* populations from our biobank (each from a different CF patient), that exhibited diversity in growth rate (Fig. 3A) and morphology (Fig. 3B). Populations from Patient 1 and Patient 2 were untypeable (UT) and did not exhibit R-pyocin killing activity (Fig. S2), while the population from Patient 3 was determined to consist of R1-pyocin producers, and did exhibit R-pyocin activity (Supplementary Dataset 2). Preliminary experimentation suggested that these populations each exhibited some susceptibility to R2-pyocins, regardless of their own R-pyocin genotype. To assess heterogeneity in susceptibility to R-pyocins within these populations, we used partially fractionated R2-pyocin cell lysate from PAO1. To show that other pyocin-associated lytic enzymes such as holin (required for R-pyocin release by forming pores in the host cell’s membrane) or lysin (murein hydrolase that degrades peptidoglycan through pores created by holin; found in an operon of several lytic genes) are not responsible for the bactericidal activity against other strains, we included lysates extracted from an isogenic PAO1 R-pyocin null mutant lacking the tail fiber gene and chaperone (PAO1ΔR). This strain has an intact lysis cassette, holin genes, and *Pf4* prophage, however filamentous prophage has previously been shown to be deactivated by chloroform (61–63), which is used during the R-pyocin isolation process to lyse the cells.

**Figure 3.**
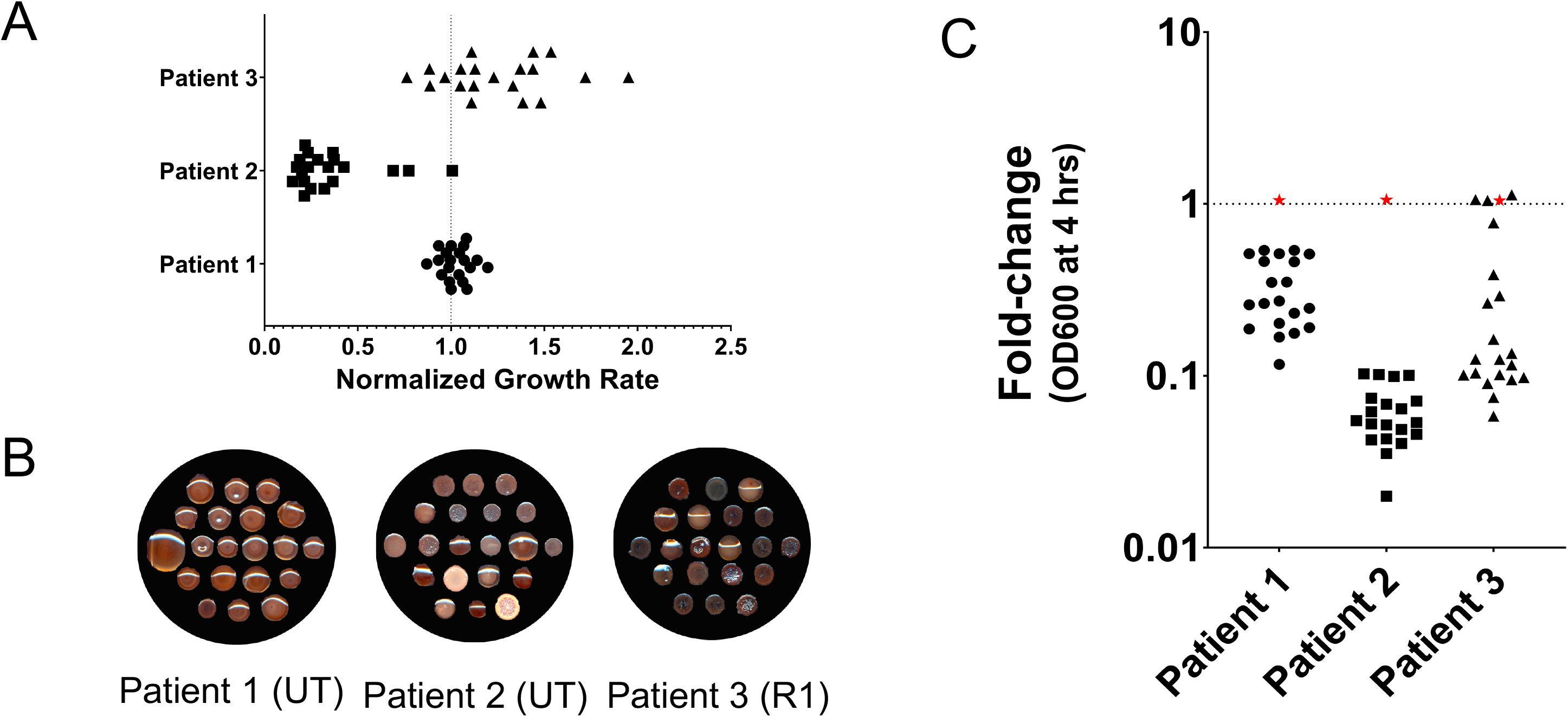
Diverse *P. aeruginosa* populations from CF exhibit heterogeneity in growth, morphology, and R2-pyocin susceptibility. (A) Growth rates vary between 20 isolates each from 3 populations collected from expectorated CF sputum. Growth rate was calculated from optical density (at 600nm) after 16 hours of growth in LB medium and normalized to PAO1. (B) The same isolates from each population are also morphologically diverse as shown on Congo Red Agar medium. Both mucoid and non-mucoid isolates are found within the same populations. The populations of Patients 1 and 2 were untypeable (UT) by classical R-pyocin typing methods, however the population of Patient 3 were typed as R1-producers. (C) The diverse isolates vary in susceptibility to the R2-pyocins of PAO1, when R2-pyocins are added to the medium during growth. Each data point represents the mean of three independent biological replicates for an individual isolate. PAO1 is denoted as a red star, and was measured with each population as a control for R2-pyocin resistance. A fold-change of 1 indicates resistance to R2-pyocin whereas a fold-change less than 1 indicates R2-pyocin susceptibility. UT = Untyped R-pyocin genotype.

By comparing growth (optical density) of *P. aeruginosa* isolates with and without R2-pyocins added to liquid culture, we found that across patients, each population as a whole exhibited different levels of susceptibility. We also found that within each population, there was heterogeneity in susceptibility among the isolates (Fig. 3C). In *P. aeruginosa* populations sourced from Patient 1 and Patient 2, all 20 isolates were susceptible to R2-pyocins, however the response varied approximately 10-fold between individual isolates. In Patient 3, the response ranged from susceptible to completely resistant. This finding demonstrates that even within one population of *P. aeruginosa* producing the same R-pyocin type, there is diversity in susceptibility to R-pyocins of other types.

### There is significant heterogeneity in R-pyocin susceptibility within a *P. aeruginosa* population

To further distinguish differences in susceptibility between isolates, we chose three isolates from the R1-pyocin producing population of Patient 3 for further analysis. We first tested the three isolates for susceptibility to R-pyocins from several R2-pyocin producing strains by visualizing killing activity with a standard, soft-agar overlay spot assay (Fig. 4A). We used R2-pyocin containing lysates from the standard laboratory strain PAO1 and a PAO1 R-pyocin null mutant lacking a functional R-pyocin (PAO1ΔR), as well as a previously described CF isolate A018 (also a R2-pyocin producer) and a corresponding R-pyocin null mutant (25, 64, 65). The spot assay indicated that Isolate 1 from Patient 3 was resistant to the R2-pyocins of PAO1 and A018, while Isolates 2 and 3 were susceptible (depicted by the zones of inhibition where the R2-pyocin lysates were spotted) (Fig. 4A).

**Figure 4.**
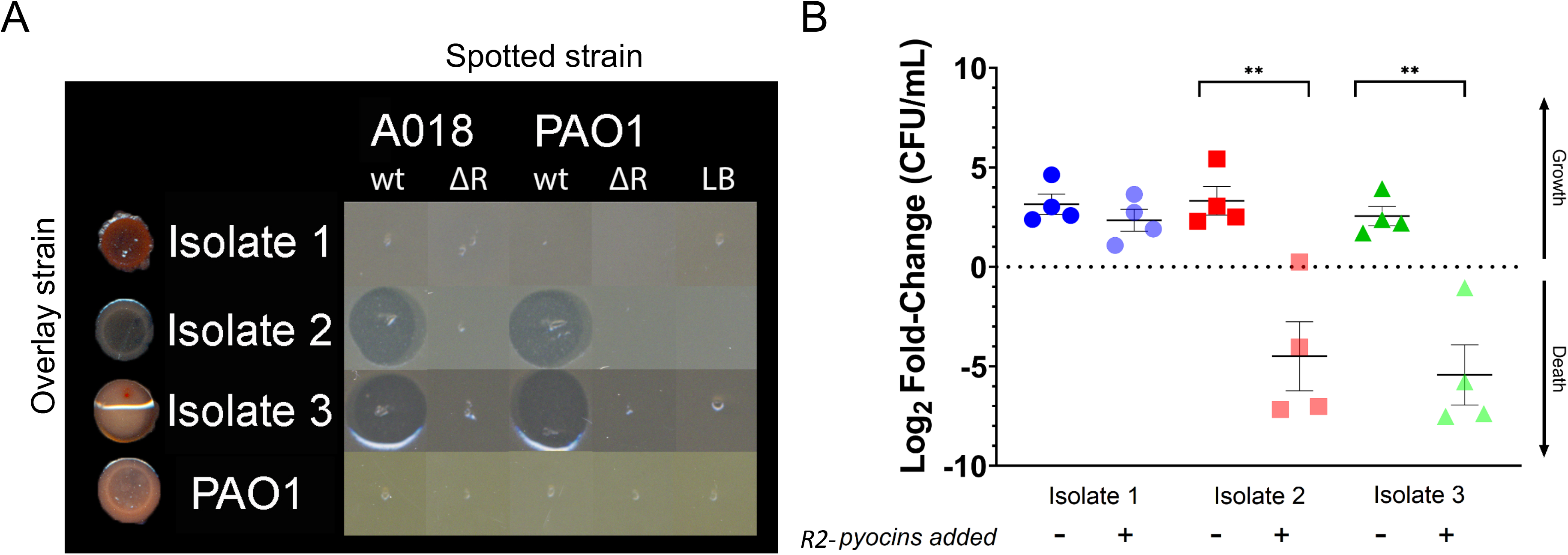
Clinical CF isolates of *P. aeruginosa* from the same population exhibit differential susceptibility to R2-pyocins. (A) Three R1-type pyocin producing clinical CF isolates from the same population (Patient 3) were tested for susceptibility to spotted lysates with a standard spot assay, and show differences in susceptibility to R2-pyocin lysates. Isolates 2 and 3 show zones of inhibition, indicating susceptibility. Spotted strains include the previously described CF strain A018 (R2), the laboratory strain PAO1 (R2), and a cell-free media preparation. (B) The same three isolates were grown in Lysogeny broth (LB), and treated with R2-pyocins extracted from PAO1. Each data point depicts a biological replicate (averaged from technical replicates). Log2 fold-change in CFU was measured at t=0 and t=4 hours for each isolate to depict growth or death in the presence or absence of R2-pyocins. Two-tailed, paired Student’s T-test between treatments confirm that Isolates 2 and 3 are susceptible to R2-pyocins (\*\**P=*0.005), while Isolate 1 is resistant (ns). A one-way ANOVA and Tukey-Kramer post-hoc test determined that Isolates 2 and 3 significantly decrease in CFU from Isolate 1 when treated with R2-pyocins (not shown, *P*<0.05).

We next measured R2-pyocin susceptibility of the three isolates with lysates from PAO1 in liquid culture, and measured Log2 fold-change of colony forming units (CFU) after 4 hours of growth to compare cell density with and without R-pyocin-containing lysates. Through this assay, we confirmed that Isolates 2 and 3 from Patient 3 are susceptible to R2-pyocins, while Isolate 1 is resistant (Fig. 4B). We confirmed in both planktonic growth and in an agar overlay that within a single population, two isolates were susceptible to R2-pyocins while one isolate was resistant, indicating that *P. aeruginosa* isolates of the same R-pyocin type and from the same population can exhibit different susceptibilities to other R-pyocins.

### R-pyocin susceptibility of isolates is dependent on the LPS core structure and not alginate production

Alginate is a type of exopolysaccharide that *P. aeruginosa* cells may secrete, giving them a characteristic mucoid morphology during growth; it contributes to a more robust biofilm and has been shown to be associated with strains that are more tolerant to antibiotics (66–68). We tested alginate production to determine whether alginate may provide resistance to R-pyocin killing or influence R-pyocin susceptibility in any way among isolates. As there was diversity in colony morphology and mucoidy among the *P. aeruginosa* population of Patient 3 (Fig. 3B), we quantified the alginate production of Isolates 1-3 from Patient 3 to determine if alginate production may influence the susceptibility differences among these isolates. We found that only Isolate 3 (R2-susceptible) produced alginate while Isolate 2 (R2-susceptible) and Isolate 1 (R2-resistant) did not produce alginate (Fig. 5A), suggesting that alginate production did not influence R2-pyocin susceptibility between the three isolates.

**Figure 5.**
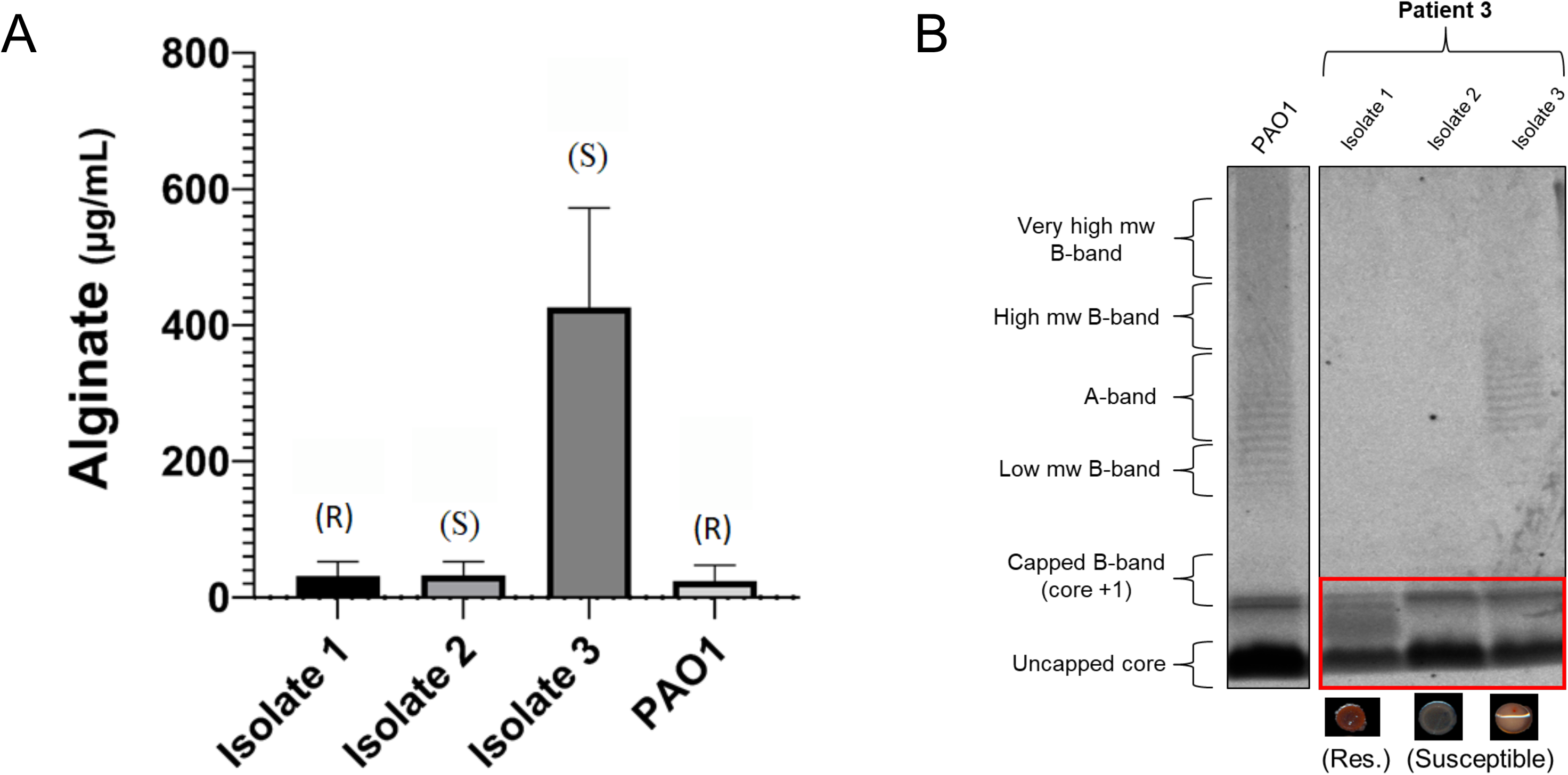
Alginate production and LPS characterization of three CF isolates of P. aeruginosa from the same population. (A) Alginate was isolated and quantified from clinical Isolates 1-3 (of Patient 3). Susceptibility to R2-pyocins is denoted by R (resistance) or S (susceptible). Alginate concentrations measured from three biological replicates are represented by the mean value and error bars depict standard error of the mean (SEM). The means of alginate concentration from each isolate were determined by comparison to a standard curve generated and analyzed with a one-way ANOVA and Tukey-Kramer post-hoc test. Isolate 3 exhibited a mucoid morphology and produces more alginate in comparison (*P*<0.025) though it is still susceptible to R2-pyocins, while Isolates 1 and 2 did not produce detectable levels of alginate (<50 μg/mL) and differ in R2-pyocin susceptibility. (B) The three isolates each exhibit different LPS phenotypes; Isolate 1 presents neither A-band nor B-band, and has a different core from PAO1. Isolate 2 also does not present A-band or B-band, but has a similar core to Isolate 3 and PAO1. Isolate 3 presents a normal core, and presents A-band antigen.

As LPS has been shown to be involved in the binding of R-pyocins to target cells (1–3, 12, 14, 15, 22–27, 29–34), we further characterized Isolates 1-3 from Patient 3 by extracting and visualizing major LPS components via gel electrophoresis and staining. All three isolates showed differences in O-specific antigen (OSA, B-band) and common antigen (CPA, A-band) presentation, but a notable difference was that the R2-resistant Isolate 1 possessed a LPS core band that migrated differently from the susceptible isolates (Fig. 5B). Along with an altered LPS core, Isolate 1 lacks both the A-band and B-band. We found that Isolate 2 lacks A-band, presented some low molecular weight B-band, and had a normal core, while Isolate 3 also had a normal core and only presented A-band. This variation in LPS phenotypes among isolates within a population is in agreement with previous studies that have shown LPS mutations frequently arise in the CF environment, both *in vivo* and *in vitro* (69–78).

### WGS sequencing of isolates reveals possible SNVs responsible for R-pyocin resistance

We obtained whole genome sequences of Isolates 1-3 from Patient 3 to determine the strain type and serotype. We found that all three isolates are of the multi-locus sequence type (MLST) 2999, suggesting they are derived from the same originating infecting strain. The serotype of each isolate was predicted to be O6, indicating that the isolates all carried the O6 gene cluster (encoding genes for synthesis of the variable O-specific antigen or B-band), and that serotype alone is likely not responsible for the differences in R-pyocin susceptibility. We found that all three isolates shared 30,127 single nucleotide variants (SNVs) when mapped against a PAO1 reference genome (Fig. 6A). We found 232 SNVs unique to the R2-resistant Isolate 1 from Patient 3, when mapped to PAO1 and compared to SNVs of the two susceptible isolates from Patient 3 (Fig. 6B). Of the 232 SNVs unique to Isolate 1, 169 SNVs were non-synonymous (Supplementary Dataset 3).

**Figure 6.**
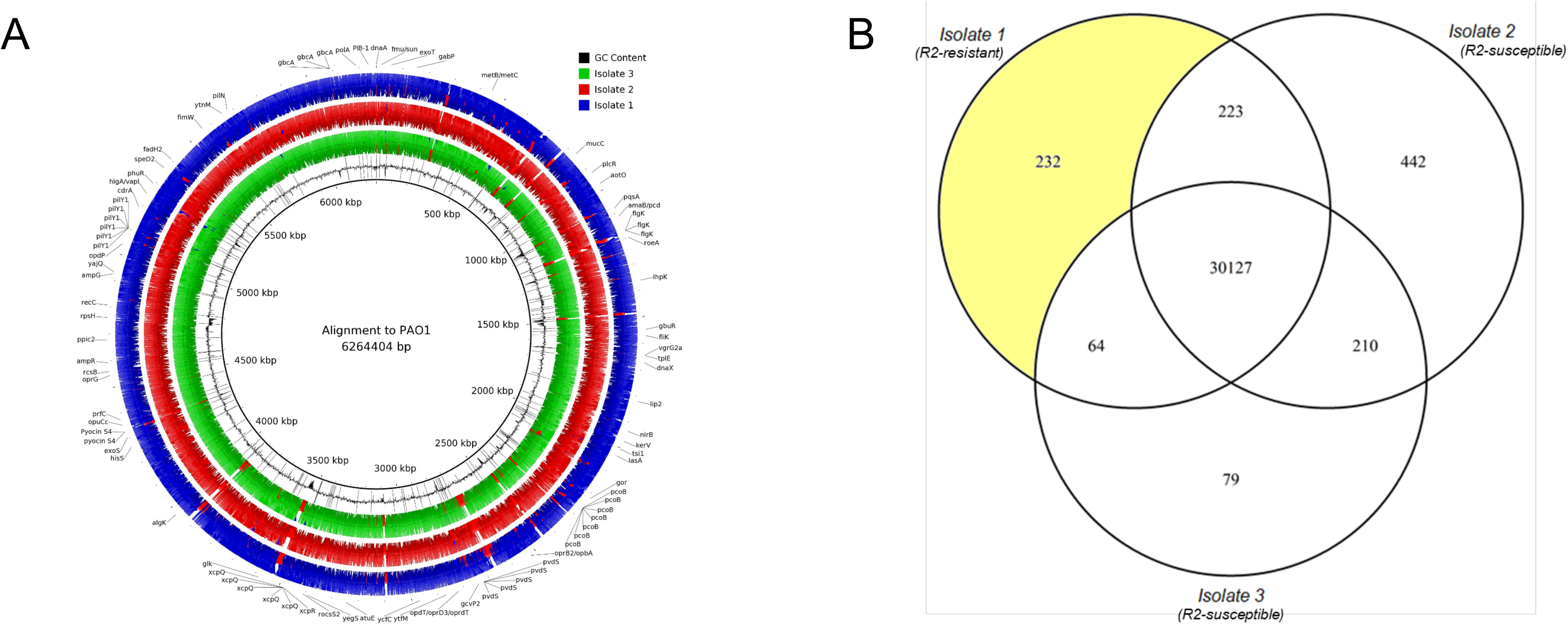
Whole-genome analysis of three CF isolates of *P. aeruginosa* from the same population. (A) Three R1-type pyocin producing clinical CF isolates from the same population were sequenced and aligned to a PAO1 reference genome for comparison. Only annotated single nucleotide variants (SNVs) unique to the R2-pyocin resistant Isolate 1 are depicted for simplicity. (B) Single nucleotide variant lists between each of the isolates when mapped to PAO1 were compared, to reveal that approximately 30,127 SNVs distinguished this population from the laboratory strain PAO1. Out of the 30,646 SNVs between Isolate 1 and PAO1, 232 were unique to Isolate 1 when compared to Isolates 2 and 3 from the same population.

Given the high number of SNVs and stark diversity between the isolates, through genetic analysis we were able to identify several candidate SNVs potentially involved in the R-pyocin resistant phenotype of this isolate, though it could be any number of these mutations in combination that play a role in the R-pyocin resistant phenotype we observed. Noteable candidate SNVs unique to Isolate 1 included a C151T (Arg51Trp) missense variant at position 833604 in the *mucC* gene (PA0765), a G710A (Arg237His) missense variant at position 1199237 of the *roeA* gene (PA1107), and a T1169C (Leu390Pro) missense variant at position 3968193 in the *algK* gene (PA3543). Each of these genes are involved in alginate production or polysaccharide biosynthesis but may indirectly influence LPS biosynthesis or presentation. However, when we complemented each of these genes (using the gene sequences from PAO1) into Isolate 1, there were no differences to R2-pyocin susceptibility; the complemented strains of Isolate 1 remained resistant to R2-pyocins (Fig. S3A) and did not alter LPS presentation (Fig. S3B).

## Discussion

Chronic infections found in CF lungs are debilitating and often lethal. *P. aeruginosa* is a common pathogen found in CF lungs and is highly resistant to multiple classes of antibiotic (4, 39–56), making the development of new and alternative therapies critical. Bacteriocins have long been considered in food preservatives as bio-controlling agents and also as therapeutics (79–85). R-pyocins have potential as an anti-pseudomonal therapeutic, but little is known about their antimicrobial efficacy against the heterogeneous *P. aeruginosa* populations found in CF lungs. We evaluated R-pyocin type and susceptibility among *P. aeruginosa* isolates sourced from CF infections and found that (i) R1-pyocins are the most prevalent R-type among respiratory infection and CF strains; (ii) a large proportion of *P. aeruginosa* strains lack R-pyocin genes entirely; (iii) isolates from *P. aeruginosa* populations collected from the same patient at a single time point have the same R-pyocin type; (iv) there is heterogeneity in susceptibility to R-pyocins within *P. aeruginosa* populations and (v) susceptibility is likely driven by diversity of LPS phenotypes within clinical populations.

We first assessed the prevalence of the different R-pyocin subtypes in *P. aeruginosa* isolates sourced from different environments by screening for R-pyocin genotypes of *P. aeruginosa* isolates through the International Pseudomonas Consortium Database (IPCD) and our biobank of CF isolates (60). We found that R1-pyocin producers make up the majority of typeable CF strains from both sources. This suggests that many CF infections may initially be colonized with R1-producing ancestor strains, or that there is a possible benefit to producing R1-type pyocins during strain competition in early stages of infection. Our findings agree with previous work, as R1-producers have been shown to be the most prevalent subtype of CF isolates evaluated in a separate study which typed 24 *P. aeruginosa* CF isolates (62.5% of which were R1-producers) (64). We found a large number of strains from the IPCD that could not be typed by our screen using classical R-pyocin typing methods, the majority of which (98%) were missing the R-pyocin structural gene cassette. Another group evaluating R-pyocin susceptibility among 34 CF isolates found that 23 (68%) of the tested isolates did not produce R-pyocins or could not be typed (28). Other work has also shown that CF isolates of *P. aeruginosa* exhibited loss of pyocin production when compared to environmental strains (85). This suggests that CF isolates of *P. aeruginosa* initially possessing R-pyocin genes may lose the ability to produce R-pyocins later in chronic infection, however further study is necessary to explore the potential fitness advantages of this evolutionary trajectory.

We are only just beginning to understand how heterogeneous *P. aeruginosa* populations impact upon treatment and disease outcomes. Previous studies have R-pyocin typed *P. aeruginosa* from CF patients and compared R-types from longitudinal collections, only evaluating single isolates from each patient (28, 64, 86). The typing of single isolates, however, does not consider the genotypic and phenotypic diversity of *P. aeruginosa* populations, which evolve over the course of chronic infection (39–49). Out of the *P. aeruginosa* populations collected from eight CF patients we found primarily R1-producing or untypeable strains. We also found that all of the isolates in each population consistently exhibited the same R-pyocin genotype over a 2-year period. A previous study typed single isolates yearly for 8 years and found that R-pyocin type could change over the course of a patient’s infection, but strain typing also showed that these patients were often infected with different strains during those events, suggesting that the observed changes in R-pyocin type were likely due to transient strains (28). Several studies have shown that diverse populations of *P. aeruginosa* from chronic infections show heterogeneity in susceptibility to a number of antibiotics, complicating susceptibility testing and treatments (39, 44, 54, 57–59). Though heterogeneity did not influence R-pyocin genotypes within our CF populations, we found that diverse populations of *P. aeruginosa* from chronic CF lung infections exhibit heterogeneity in susceptibility to R-pyocins. Specifically we found that (i) each population of *P. aeruginosa* as a whole exhibited differential levels of susceptibility and (ii) within each population, there was heterogeneity in susceptibility among individual isolates.

Previous studies have shown the effect of various LPS mutations on R-pyocin susceptibility in laboratory strains PAO1 and PAK (24–26), however, to our knowledge, our work is the first to examine LPS phenotypes associated with R-pyocin susceptibility and resistance among clinical isolates. We found that differences in the core of the LPS appear to be correlated with the differences in susceptibility and resistance to R-pyocins among three isolates from one patient; specifically, that a resistant CF isolate produces an altered LPS core. Our findings are in agreement with recent work demonstrating heterogenous resistance mechanisms and persistence against tailocin killing by alterations of the LPS and associated glycosyltransferases in *Pseudomonas syringae* (87). It is unknown if the altered LPS core phenotype we have found is commonly identified in CF *P. aeruginosa* populations, or if there is a fitness benefit it confers during infection. During initial colonization and strain establishment in the early stages of infection, the LPS (and therefore R-pyocin susceptibility) may play a role in antagonistic interactions between competing strains and ultimately determine the dominating strain.

While investigating possible mechanisms for the variable R2-pyocin susceptibility of Isolates 1-3 from Patient 3, we showed that serotype, strain type, and alginate production were not responsible for the resistance in Isolate 1. When we probed the genomes of three isolates from a single clinical population exhibiting heterogenous susceptibility to R2-pyocins, we identified several candidate SNVs in *mucC* (PA0765), *roeA* (PA1107), and *algK* (PA3543), unique to the resistant isolate; however ultimately these genes did not effect R2-pyocin resistance or LPS presentation when complemented. All three genes are involved with the alginate production pathway or other extracellular polysaccharides (41, 88–91), known to be closely intertwined with various LPS synthesis and secretory pathways; thus it is possible that any combination or all three variants combined may play a role in indirectly impeding presentation of various portions of the LPS or core residues. As there are many regulatory mechanisms and biosynthetic genes still yet to be understood that are involved in LPS synthesis and presentation, it is also possible that there is a mechanism at play that has yet to be described.

Overall our findings highlight that clinical populations of *P. aeruginosa* exhibit a heterogenous response to R-pyocins and that this likely extends to any antimicrobial that utilizes the LPS. A number of LPS- and saccharide-binding phage have been identified and tested as antimicrobials against *P. aeruginosa* (24, 102–108) and other types of LPS-specific antimicrobials, including R-pyocins, are also being considered as treatments (12, 13, 37, 38, 109, 110). Our work implies that treatment with alternative therapies utilizing the LPS may not eradicate strains within infections completely, potentially leading to highly resistant isolates taking over. This reiterates the importance of assessing multiple, diverse isolates from populations of *P. aeruginosa* rather than taking a single isolate from a population. To better understand how R-pyocins and other LPS-binding antimicrobial therapies can be utilized for alternative treatments, it is crucial to know more about the LPS, core phenotypes, and how they are evolving in *P. aeruginosa* populations during infection, to better understand the resulting impact on the potential efficacy of these antimicrobials.

## Materials and Methods

### Bacterial strains, media, and culture conditions

Expectorated sputum samples for this study were collected from adult CF patients through Emory–Children’s Center for Cystic Fibrosis and Airways Disease Research by the Cystic Fibrosis Biospecimen Laboratory with IRB approval (Georgia Tech approval H18220). *P. aeruginosa* populations were collected from each sputum sample using selective media (Pseudomonas Isolation Agar, Sigma-Aldrich) before isolating single colonies for further characterization. In total, 204 isolates from our biobank, sourced from CF chronic lung infections, were studied for R-pyocin typing, and 60 of these isolates were chosen at random for further study (20 isolates from three different patients). All *P. aeruginosa* isolates, laboratory and indicator strains used in this work are listed and described in Supplementary Dataset 2. All bacterial cultures were grown in lysogeny broth (LB) medium at 37°C with 200rpm shaking agitation. Standard genetic techniques were used for construction of *P. aeruginosa* mutants. The construction of plasmids and generation of mutants can be found in more detail in Text S1 of the supplemental material. All plasmids and primers used for mutant generation can be found in Table S1.

### Assessing colony morphology diversity in clinical isolates

To evaluate the diversity in colony morphologies between *P. aeruginosa* clinical isolates, we used a Congo Red-based agar media (1% agar, 1×M63 salts (3 g of monobasic KHPO4, 7 g of K2PO4, and 2 g of NH4.2SO4, pH adjusted to 7.4), 2.5 mM magnesium chloride, 0.4 mM calcium chloride, 0.1% casamino acids, 0.1% yeast extracts, 40 mg/l Congo red solution, 100 μM ferrous ammonium sulfate, and 0.4% glycerol) (111). Isolates were grown overnight (16-18 hours) in LB medium at 37°C with 200rpm shaking agitation before spotting 10 μL onto the plates. We incubated the plates overnight at 37°C, and for a further 4 days at 22°C. The colonies were imaged with an Epson scanner at 800 dpi.

### R-pyocin typing strains of the IPCD

Previously characterized *P. aeruginosa* strains were selected for query sequences for each R-type: PAK an R1, PAO1 an R2, and E429 an R5 (25). As the R-pyocin tail fiber sequences of PAO1 (NP_249311.1) (112–114) and PAK (Y880_RS29810; accession GCA_000568855.2) (115) are annotated, the corresponding nucleotide sequences were downloaded in FASTA format from NCBI. The E429 (R5) tail fiber sequence was identified from previous work (25) and downloaded from NCBI by using the PAO1 tail fiber sequence (PA0620) to BLAST against the E429 genome (60). NCBI’s BLAST+ was also used to determine percent identity between the whole tail fiber sequences of each strain/R-type pair-wise, in order to determine the variable region of the tail fiber gene to use for typing. Homology information, percent identity, query coverage and parameters can be found in Supplementary Data 1 (116). A detailed description of R-pyocin typing the IPCD can be found in Text S1 in the supplemental material.

### PCR conditions and R-pyocin typing

A detailed description of R-pyocin typing the biobank of CF populations can be found in Text S1 of the supplemental material. R-pyocin typing primer details and product sizes are listed in Table S1.

### Expression and extraction of R-pyocins

Expression of R2-pyocins used in the susceptibility assays were induced and pyocins extracted from PAO1 with a modified method from published work (12) and can be found in detail in text S1 of the supplemental material.

### Spot assay for R-pyocin activity

Overnight cultures of clinical *P. aeruginosa* isolates were used to inoculate 4 mL of cooled soft top (0.7%) agar at an optical density at 600 nm (OD_600nm_) of 0.01. For complemented strains, 4μL of Chromomax isopropyl β-D-1-thiogalacto-pyranoside (IPTG)/X-Gal solution (Fisher Scientific) was added per 1mL of soft agar for induction of gene expression. This mixture was poured onto LB agar plates for an overlay of the indicator strain. For Fig. 4A, R2-pyocin lysates extracted from PAO1, A018, and respective R-pyocin null mutants and were vortexed before spotting 5 μL of each lysate onto the soft-top overlay indicator strains. For Fig. S2, lysates were collected from clinical isolates as described in text S1, were vortexed and serially diluted in LB 10-fold before spotting 5 μL of each dilution onto the soft-top overlay of previously described indicator strain A026 (64). Spots were allowed to dry before plates were incubated at 37°C overnight. Clear zones of growth inhibition indicated R-pyocin dependent activity against the indicator strains. Spot assays for R-pyocin susceptibility were conducted in triplicate.

### Microtiter plate method for R-pyocin activity

R2-pyocin susceptibility was measured by changes optical density under conditions with and without added R2-pyocins, using a 96-well microtiter plate. A detailed description of the microtiter plate method can be found in Text S1 of the supplemental material.

### Alginate Isolation and Quantification

Alginate was isolated and quantified by carbazole methods described previously (117, 118) and described in detail in Text S1 of the supplemental material.

### Statistical analysis

Data analysis was performed using Prism (Graphpad software, version 9). Population heterogeneity experiments were normalized to a corresponding R-pyocin null mutant lysate control culture for each isolate (to account for heterogeneity in growth rates) for fold-change, and analyzed with Prism. CFU data was analyzed for four isolates by normalizing CFU measurements at t=0 hours and t=4 hours for cultures with and without R-pyocin lysates to compare log_2_ fold-change between treatments for each isolate. Statistical significance between treatments for each isolate was measured with a paired, parametric Student’s T-test. Differences across all three isolates with the R-pyocin treatments and alginate production were analyzed using a one-way analysis of variance (ANOVA) with Tukey’s multiple comparisons analysis in Prism. Fig. 3C and Fig. 5A show mean values of three biological replicates for each individual isolate, while Fig. 4B shows four biological replicates (averaged from technical replicates) as individual data points. Error bars represent standard error (SEM) and significance is denoted as follows: **, *P =* 0.005.

### LPS extraction and characterization

LPS of bacterial cultures were isolated and visualized as described by previous methods (119) and described in detail in Text S1 of the supplemental material.

### Whole genome sequencing analysis

A detailed description of genomic DNA preparation, sequencing, and analysis can be found in Text S1 of the supplementary material. The full list of SNVs unique to each isolate can be found in Supplementary Data 3.

### Data Availability

All sequences were deposited in the National Center for Biotechnology Information’s SRA database under the accession number PRJNA67996 (120). Raw data and code have been made available in the Dryad Digital Repository: https://doi.org/10.5061/dryad.573n5tb67 (121).

## Supplemental Material

Table S1. Primers and plasmids used in this study.

Supplementary Dataset 1. Detailed description of bacterial strains used in this study. Supplementary Dataset 2. Detailed results of R-pyocin typed IPCD strains.

Supplementary Dataset 3. Detailed description of whole genome sequencing parameters and SNV lists.

Text S1. Detailed description of materials and methods.

Text S2. Supplemental references.

## Acknowledgements

We thank Freya Harrison and Joanna Goldberg for comments on the manuscript. We thank Arlene Stecenko and the Cystic Fibrosis Biospecimen Registry (CFBR), maintained by Emory University and the Children’s Healthcare of Atlanta Center for Cystic Fibrosis and Airways Disease Research (CF-AIR), for clinical CF populations of *P. aeruginosa*. This work was supported by the Georgia Institute of Technology and The Cystic Fibrosis Foundation for a Pilot Grant (DIGGLE18I0) and a Research Grant (DIGGLE20G0) to S.P.D. M.M., J.T. and S.P.D. conceived the study and wrote the manuscript. J.T. generated the PAO1ΔR mutant strain. M.M. performed the experimental work and analyzed data.

---

**Figure S1.**
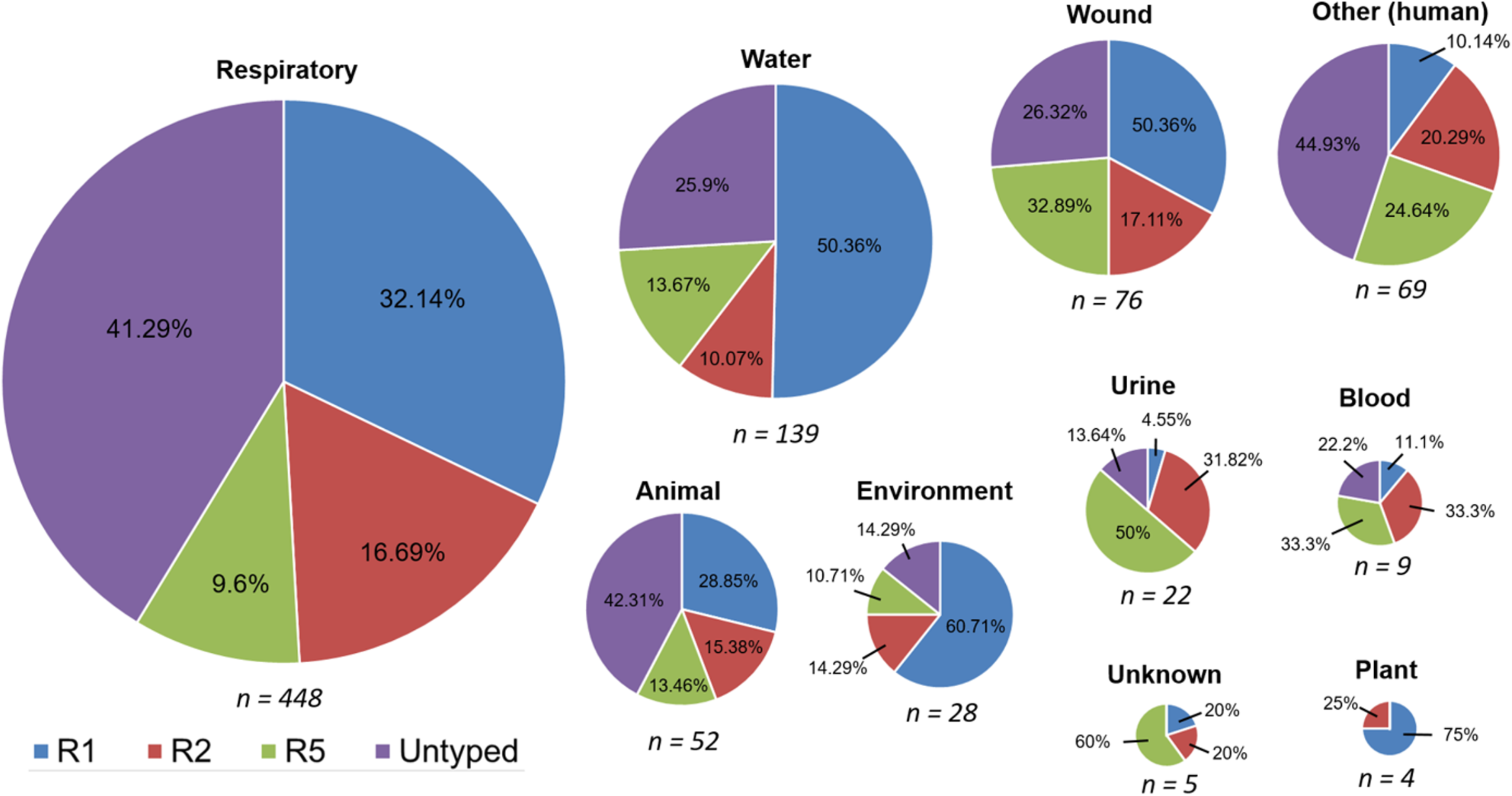
Distribution of R-pyocin types across *P. aeruginosa* strains isolated from various sources as a part of the IPCD. Using BLAST+, 852 publicly available strains through the International Pseudomonas Consortium Database (IPCD) were assessed for R-pyocin genotype. Of the 852 strains, 549 strains were typeable (assigned to either R1 - R2- or R5-pyocin types). The distribution of R-pyocin types among typeable respiratory isolates suggest that R1-pyocin producers are the most prevalent subtype found, including in CF and in other types of respiratory infections.

**Figure S2.**
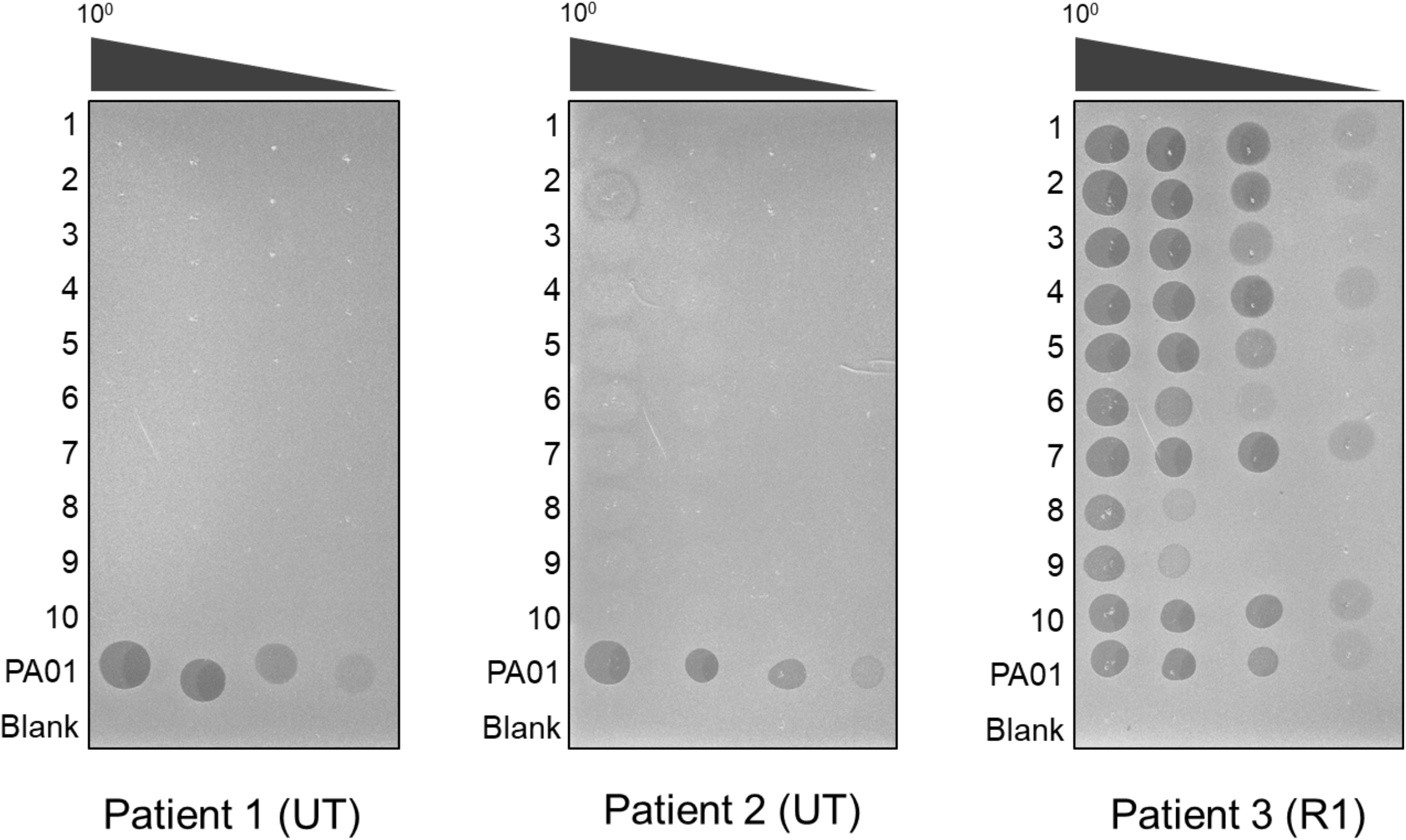
Untypeable *P. aeruginosa* isolates from two CF patients do not exhibit R-pyocin activity. From each population of Patient 1, Patient 2 and Patient 3, isolates 1-10 were tested for R-pyocin activity. Partially fractionated lysates were serially diluted 10-fold in lysogeny broth (LB) and each dilution was spotted (5 μL) onto an agar overlay inoculated with a previously described CF strain, A026. Untypeable (UT) isolates from Patients 1 and 2 do not exhibit R-pyocin activity, while isoaltes from Patient 3 (R1-pyocin genotype) does show killing activity agains the indicator strain. PAO1 (an R2-pyocin producer) is included as a control for killing activity. Zones of inhibition indicate R-pyocin killing.

**Figure S3.**
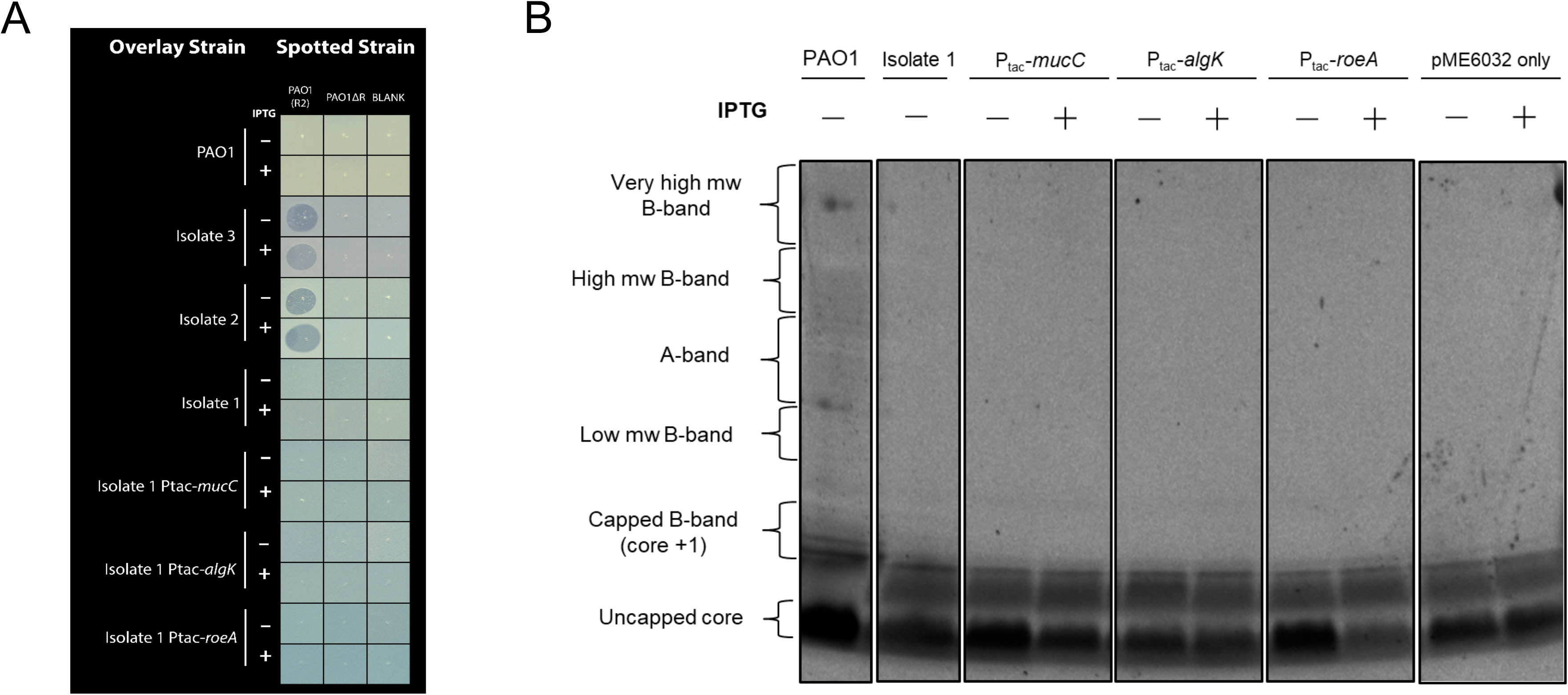
R2-pyocin resistance and LPS presentation in complemented strains of clinical Isolate 1 from Patient 3. (A) The clinical Isolates 1-3 from Patient 3 depicted in a spot assay with strains of Isolate 1 complemented with *mucC*, *algK*, and *roeA* of PAO1 inducible under P_tac_. Isolates 2 and 3 show susceptibility to the R2-pyocins of PAO1 (denoted by the zones of inhibition), whereas Isolate 1 and all complemented strains of Isolate 1 are resistant. Each isolate or strain was tested with and without Chromomax IPTG/X-Gal for induction. (B) The complemented strains of Isolate 1 were extracted and visualized by SDS-Page, confirming no alterations to the LPS presentation resulting from complementation. LPS extractions without the Chromomax induction agent were included.

